# Segregation-to-Integration Transformation Model of Memory Evolution

**DOI:** 10.1101/2023.12.05.570097

**Authors:** Luz Bavassi, Lluís Fuentemilla

## Abstract

Memories are thought to use coding schemes that dynamically adjust their representational structure to maximize both persistence and utility. However, the nature of these coding scheme adjustments and their impact on the temporal evolution of memory after initial encoding is unclear. Here we introduce the Segregation-to-Integration Transformation (SIT) Model, a neural network formalization that offers a unified account of how the representational structure of a memory is transformed over time. SIT Model asserts that memories initially adopt a highly modular or segregated network structure, functioning as an optimal storage buffer by striking a balance between protection from disruptions and accommodating substantial information. Over time, a repeated combination of neural network reactivations, spreading, and synaptic plasticity transforms the initial modular memory structure into an integrated memory form, facilitating intercommunity spreading and fostering generalization. In addition, SIT Model reveals the existence of an optimal window during this transformation where memories are most susceptible to malleability, suggesting a non-linear or inverted U-shaped function in memory evolution. The results of our model integrate a wide range of experimental phenomena along with accounts of memory consolidation and reconsolidation, offering a unique perspective on memory evolution by leveraging simple architectural neural network property rules.

## 1 Introduction

Memories, akin to other natural systems, evolve to enhance survival within organisms. Their perpetuity is constrained by evolving experiential needs, prompting memories to adapt the coding scheme of their representational structures for optimized persistence and efficacy. Despite consensus on memory evolution entailing architectural changes, understanding the balance between memory persistence and efficacy remains limited. A key research challenge involves elucidating how experience-induced efficacy influences memory changes, promoting persistence.

Our current understanding of memory evolution suggests that initially, memories are encoded by groups of neurons with synchronized activity, forming clustered neural ensembles, that if reactivated, induce memory retrieval [Josselyn and Tonegawa, 2020]. These neural ensembles are identified in the hippocampus, but over time, they undergo structural changes, losing their modular properties in the hippocampus [Gonzalez et al., 2019], and other neural ensembles beyond the hippocampus, such as the prefrontal cortex, gradually assume their representation [Frankland and Bontempi, 2005]. This transition, however, occurs relatively slowly and requires repeated reactivation of the initial neural ensembles to shift the structure of the neural ensembles between networked regions [Frankland and Bontempi, 2005]. For a while, the dominating view was that once memories shifted to neocortical neural ensembles, they became consolidated and stable in the long term. More contemporary views advocate for the notion that consolidation and reactivation may act in the service of generalization [Sun et al., 2023]. That is, given that individual memorized experiences rarely repeat exactly, generalization allows us to identify systematic relationships between features of the world, ultimately involving extending learned information to novel contexts. Thus, generalization involves the process of linking and extracting commonalities among various memories. As the brain generalizes, initial memory representations undergo a transformative shift in their structure, becoming intricately connected to related memories [Nadel and Moscovitch, 1997, Winocur et al., 2010]. Through this interconnected web of associations, the initially clustered memory representations become part of a broader network, where each memory influences and is influenced by others. These processes ensure that consolidated memories are not isolated but integrated into an interconnected form, making them applicable to a range of situations and contributing to the adaptive evolution of memories.

The idea that consolidated memories are not static but undergo continual evolution is also supported by research indicating that reactivating seemingly stable memories can make them temporarily labile and susceptible to reconsolidation [Eisenberg and Dudai, 2004, Hardt et al., 2010, Nader et al., 2000, Nader, 2003, Sara, 2000]. However, memory reconsolidation dynamics also change with memory age; young memories are susceptible to changes via reactivation, while older ones are less malleable and more resistant to modification [Alberini, 2011, Milekic and Alberini, 2002, Inda et al., 2011, Forcato et al., 2013, Fernández et al., 2016], indicating the existence of an optimal memory malleability window for memory reconsolidation. While the neural mechanisms supporting the existence of an optimal window are still not clearly understood, it is assumed that reconsolidation occurs if new memories trigger the reactivation of the hippocampal networks that were active during original learning [Debiec et al., 2002, Hupbach et al., 2007, Winocur et al., 2009]. On the other hand, memories that have already shifted to a generalized network form would be minimally affected by reactivation [Dudai, 1996, McKenzie and Eichenbaum, 2011]. These findings suggest that memory evolution requires a deeper understanding of the relationship between memory reactivation and representational transformation over time.

We propose SIT: the Segregation-to-Integration Transformation model, a unified account of how the representational structure of a memory is transformed via repeated reactivations over time. SIT builds up on the notion that memories can be represented as neural network ensembles, including a collection of neurons (nodes) linked by pairwise connections (edges), and that memory transformation is accounted by changes in the connectivity pattern between network ensembles [Ryan et al., 2015, 2021a, Tonegawa et al., 2015, Ortega-de San Luis et al., 2023]. Our goal was to develop a simplified model of memory evolution that captured the properties of memory transformation outlined by models of memory consolidation and reconsolidation, Figure 1. However, unlike dual-representational models involving the differential contribution of the hippocampus and neocortex [Sun et al., 2023, McClelland et al., 1995], we aimed to examine whether memory transformation can be explained by principles of information spread in a one-layer network structure, that has been effective to capture recurrent information transmission and retention in other complex network systems, such as in social networks [Centola, 2010, Danon et al., 2008, Newman, 2006, Weng et al., 2013].

**Figure 1.**
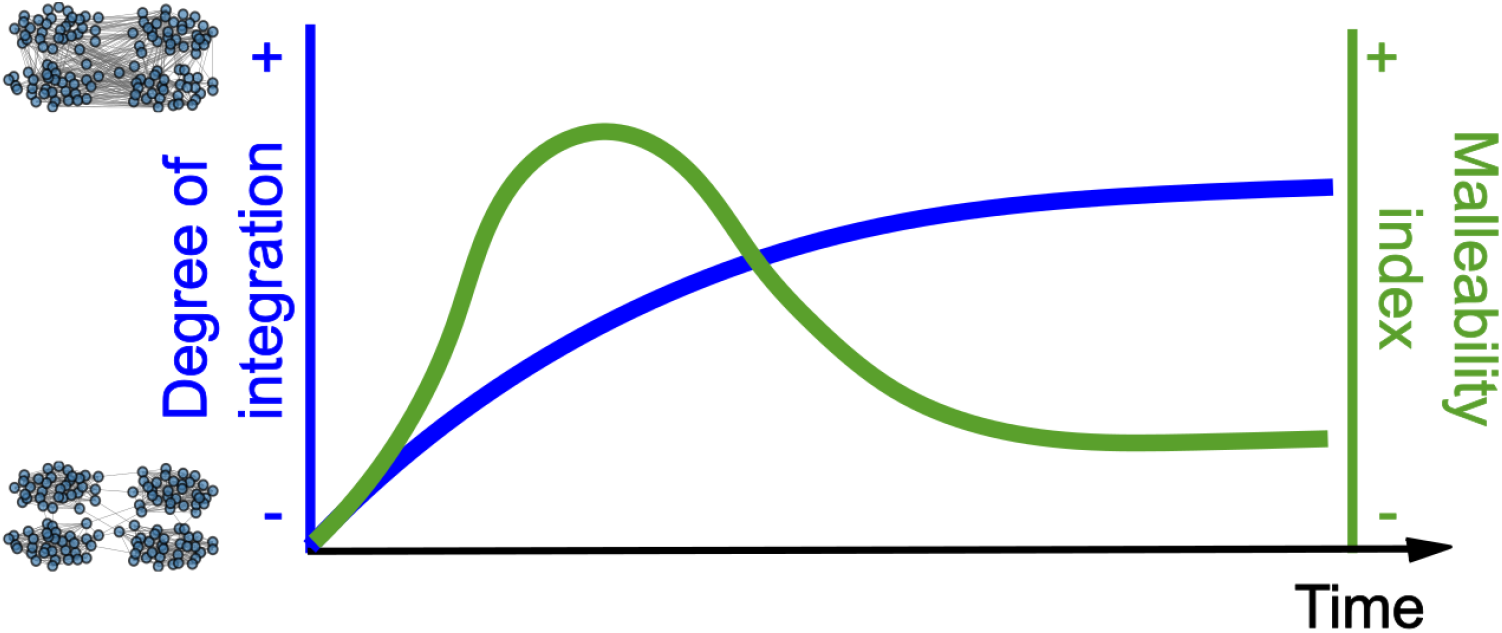
Schematic depiction of the Segregation-to-Integration Transformation (SIT) Model. Memory networks shift from a segregated into an integrated form over time, blue line. The model postulates the existence of an optimal memory malleability window in an early stage, green line.

The central tenet of our proposal is that memories tend to shift from highly segregated (modular) to intricately integrated network forms and that this process is guided by a combination of repeated network reactivations, spread of activation, and plasticity rules. We hypothesized that memories are initially stored as highly modular networks, as they serve as a buffer for activation spread within the community where they originated rather than spreading throughout other community structures in a network [Newman, 2006]. Additionally, a sparsely segregated network expands the range of possible configurations, thereby enhancing storage capacity [Brunel, 2016]. Consequently, a segregated network organization represents an optimized configuration for early memories, effectively balancing the need to shield the network from disruptions or interference while accommodating a substantial amount of information storage. Over time, repeated reactivation of elements of the original network would transform the highly modular network structure into a more integrated form, integrating links to nodes representing new overlapping information. The resulting network configuration, characterized by a low modular structure, would help intercommunity spreading, thereby promoting generalization. A crucial insight from our modeling approach is the finding of an optimal window during this transformation whereby memories are most susceptible to malleability. This implies that the process of memory evolution follows a nonlinear or inverted “U-shaped” function, thereby highlighting that memories are ever-changing but susceptible to being optimally malleable at specific stages of the transformation process. The resulting model not only unifies various experimental phenomena but also integrates memory consolidation and reconsolidation accounts, offering a perspective by which simple architectural neural network property rules can account for memory evolution.

## 2 Results

Our goal was to study the dynamics of memories transformation over time. We conceptualized memories for events as neuronal ensembles distributed throughout the brain, rather than being confined to particular regions [Vetere et al., 2017, Roy et al., 2022, Ryan et al., 2021a] that are susceptible to being reactivated spontaneously or cued by elements of the experience that partially overlap with the original event. We hypothesized the repeated reactivation of elements of the original neuronal ensemble would induce structural changes to the network.

To systematically investigate the changes in this evolving process, we prepared an ensemble of networks with four communities with an equal number of nodes and connecting links within communities. In our model, each node could achieve one of two possible values: *s* = {0, 1}, where 1 represented the “active” mode and 0 was the “inactive” one. Unweighted undirected edges illustrated the structural pattern of connections between nodes (1 for nodes that were co-active and 0 for nodes that were not co-active). We studied the dynamics of memory transformation by modeling network changes subserved by creating and removing edges following a Hebbian plasticity rule. As neural plasticity can be extended to connected neurons not directly activated [McKenzie et al., 2021, Mugnaini et al., 2023], our model incorporated a deterministic spreading rule throughout the network, enabling the influence of activation beyond the initially active neurons.

We formulated our model’s findings to illustrate different states in the progression of memory. We borrowed the terminology associated with the classification of star evolution to designate each of them in our model. First, we characterized the initial state of memory as a set of nodes exhibiting a highly modular structure (*“Radiant stage”*: Early memory formations). Following this, we simulated the network-level consequences ensuing from a single reactivation (*“Celestial stage”*: Memory evolution). Finally, we investigated the network reconfiguration effects of repeated reactivations over time (*“Enlightened Stage”*: The ultimate state), Figure 2.

**Figure 2.**
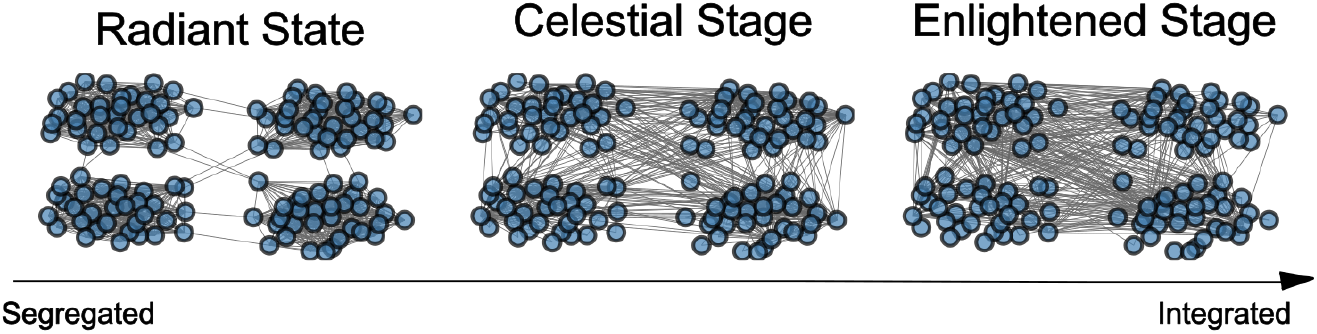
Examples of three networks with different degrees of integration (*Z*). All networks have the same number of nodes (*N* = 128) and they were built by joining four similar communities of 32 nodes each.

### 2.1 Memory stages

#### 2.1.1 “Radiant stage”: early memory forms

We aimed to develop a low-dimensional model that captured the fundamental properties described in extant models of memory consolidation and reconsolidation. In these models, memories are initially encoded in a segregated form, described by neuron ensembles with highly synchronized activity. This co-firing pattern of activity describes jointly active neurons organized into motifs, or groups of neurons of high modular activity within a larger network, that underpin spatially selective assemblies representing memories for experienced events [O’Neill et al., 2008]. In our model, we defined the first state of memory as a network model with four communities, each of them with the same number of nodes and connections. This segregated form means that nodes can be split into internally dense and externally sparse community subnetworks.

#### 2.1.2 “Celestial stage”: memories evolve

Rodent studies have demonstrated that stable synchronous activity in initially segregated neuronal ensembles can contribute to memory persistence [Gonzalez et al., 2019]. However, the process of a single memory reactivation induces topological reorganization in the co-activity structure of the network [Gava et al., 2021]. Then, we aimed to characterize the impact of this single memory reactivation on the topographical arrangement of the initial memory form. To computationally simulate this phenomenon, we employed a reactivation strategy involving the activation of a random set of nodes and examined the ensuing propagation of this activity the network. We characterized this spreading activity with a deterministic rule, assuming that reactivation reach more features of a memory than the ones that are directly represented in the original neural ensemble [Anderson, 1983]. We modeled this cascade dynamics in three steps: Turn-on/Activation, Spreading, and Plasticity defined as follows:

##### Turn-on

We represented a memory by a collection of *N* nodes linked by *M* undirected edges. Nodes can either be in active (1) or in an inactive (0) state. An input that overlaps with an original experience activate a collection of nodes of the original memory transitioning its mode from an inactive to an active state (from 0 →1). We quantified the fraction of active nodes under the term “Intensity” (*Int*). We assumed that the larger the degree of overlap between a current input and an existing memory would increase *Int* of the resulting network, and we expected that coactivation of nodes would lead to changes in the connections between these neurons [Ritvo et al., 2019, Wammes et al., 2022]. We simulated the impact of reactivation covering a range of possible nodes turned active, ranging from 0.1 to 0.6, assuming that larger *Int* values would reflect greater overlap between original and reactivated memory.

##### Spreading

We adopted a linear threshold model, commonly applied in studies of information diffusion in social networks [Nematzadeh et al., 2014, Weng et al., 2013, Centola, 2010], to account for activity propagation in the network. We formally defined the linear threshold model as follows. As explained before, the state of a node *i* at time *t* was expressed by a binary variable *s*_*i*_(*t*) = { 0, 1}, where 1 denoted an “active” state and 0 an “inactive” one. At time *t* = 0, a fraction of randomly chosen nodes were initialized in the active state. In the subsequent step, the state of each node was updated according to the following threshold rule, Equation 1:

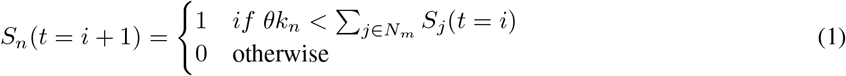

where *θ* was the threshold parameter, *k*_*n*_ was the node degree of node *n* and *j* ∈ *N*_*m*_ represented the set of neighbors of node *n*. The spreading persisted until the network reached a stable state, defined as *S*_*n*_(*t* = *i* + 1) = *S*_*n*_(*t* = *i*) ∀*n*. We conceived the threshold parameter (*θ*) as a proxy of neuronal excitability state of a network nodes. In our modeling, we systematically varied a the values to assess for the possibility, aligning with rodent literature, that fluctuations in neuronal excitability determine how memory ensembles interact, promoting either memory integration or separation [Josselyn and Frankland, 2018a].

##### Plasticity

Once the network reached a stable state, a *Hebbian* plasticity rule was implemented. This step reconfigured the connectivity matrix of the neural network by creating an edge between two nodes *L*_*n,m*_ if both connected nodes were in an active mode and removing it if both two connected nodes were in an inactive mode, Equation 2:

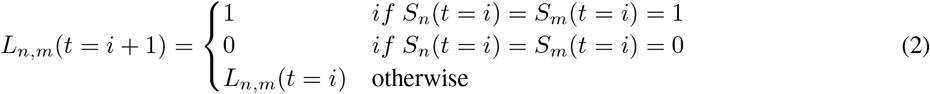

##### Degree of integration (*Z*)

We hypothesized that memories would tend to shift from highly segregated to richly integrated state forms. To formalize the state of the network, we quantified the degree of integration following Equation 3:

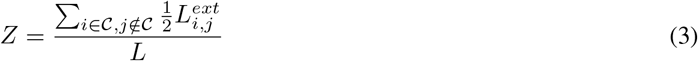

where *L* is the total number of edges and 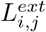 are edges that linked node *i*, a member of the initial community 𝒞, with node *j* that belongs to another community. Thus, *Z∼* 0 would define a highly segregated network, with dense intracommunity connectivity pattern and few intercommunity links (Figure 2 - left). With an increasing number of edges between communities, the level of integration, represented by the parameter *Z*, rises, eventually reaching a value of *Z∼*1 (Figure 2 - right).

To systematically investigate the impact of a single reactivation in an early-form network, we quantified changes in the connectivity pattern with varying intensities (*Int*) and excitability thresholds (*θ*). Specifically, we focus on two measures: changes in the degree of integration (Δ*Z*) and changes in the number of connections. For this, we defined the malleability index (Δ*L*) as the number of created and removed edges relative to the total number of edges in the network.

First, we examined how a single reactivation produces changes in the degree of memory integration. Figure 3A) depicts the transition in the degree of integration after reactivation (*Z*_*f*_) by varying the levels of *Int* as a function of the initial state (*Z*_*o*_), for fixed *θ*. We found that reactivation with higher levels of *Int* always jumped to a fully integrated form. In addition, reactivation with lower levels of *Int* linearly increased the degree of integration. Notably, *Z*_*f*_ consistently exceeded *Z*_*o*_ for every level of *Int*, suggesting that even a single reactivation can transform the initial network state into a more integrated form. This point holds significant importance because although the SIT model suggested a unidirectional shift, it was not inherently imposed in the reactivation rule itself; rather, it emerged because of the spreading of activation and the application of the plasticity rule. In addition, we analyzed the role of the excitability threshold (*θ*) in shifts of the degree of integration (Δ*Z*, Equation 4), as shown in Figure 3B. Interestingly, shifts of the integration state of the network produced by the reactivation depend weakly on *θ*, Δ*Z* appears almost invariant to *θ* fpr a fixed *Int* = 0.3.

**Figure 3.**
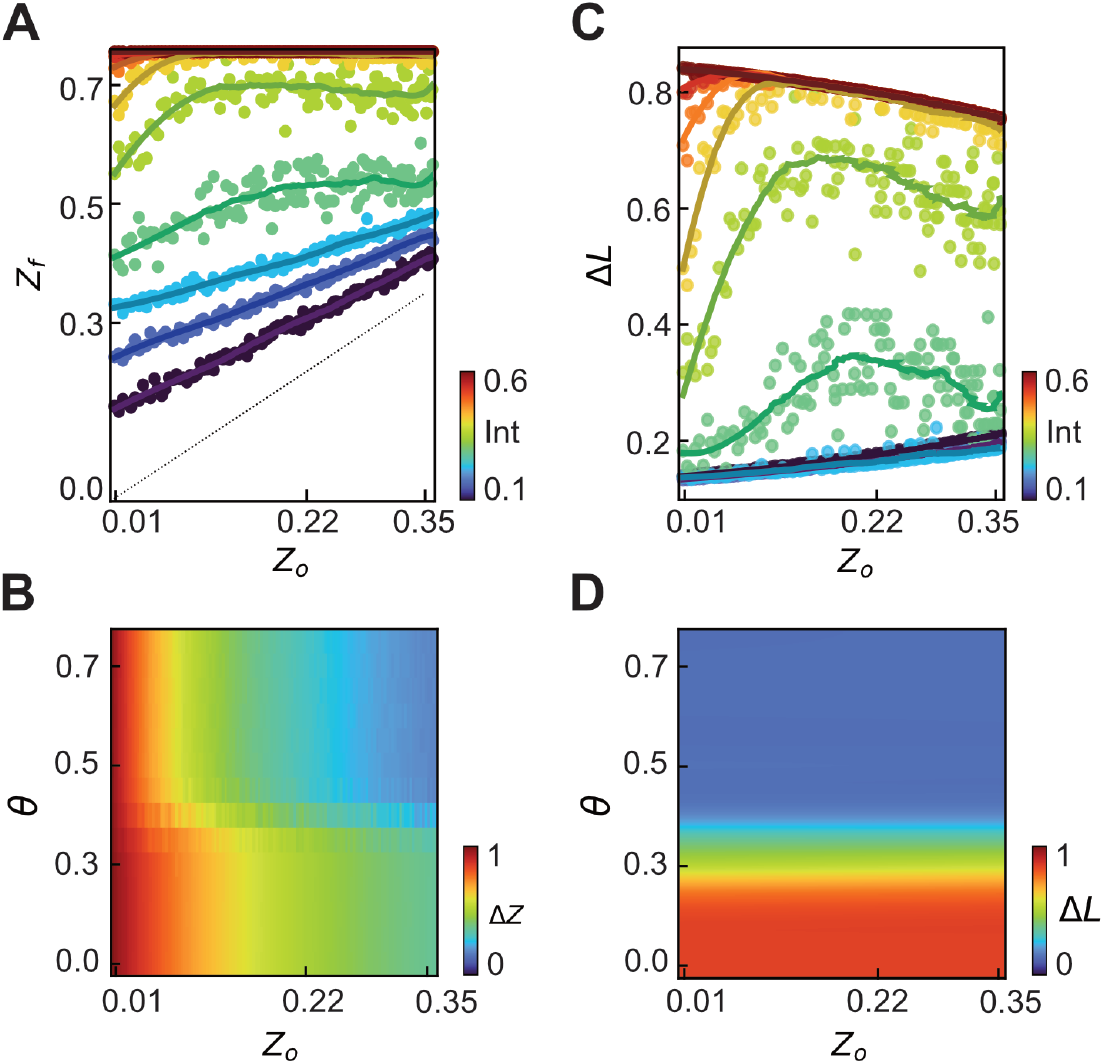
Description of reactivation concerning Intensity, the degree of integration, and the excitability threshold. **A**. Transition from *Z*_*o*_ → *Z*_*f*_ after a reactivation with *θ* = 0.4. Dots depict the relation between the degree of segregation pre (*Z*_*o*_) and post (*Z*_*f*_) reactivation. Cold colors represent reactivations with low intensities while warm colors represent higher intensities. **B**. Heatmap of Δ*Z* as a function of the degree of integration of the former network and threshold of propagation. Here, intensity is fixed (*Int* = 0.3). **C**. Relation between the malleability index and the degree of integration of the former network, *θ* = 0.4. Colors represent the reactivation *intensity*. **D**. Heatmap of the malleability index as a function of the degree of integration of the former network and excitability threshold. *Intensity* is fixed (*Int* = 0.3). The reported data is an average of multiple experimental runs (25) where the activation of nodes (turn-on) was randomized in each run.

Next, we addressed the changes in the number of connections by quantifying the Malleability Index (Δ*L*, Equation 5). Δ*L* exhibited a pronounced dependence on *Int*, Figure 3C. A lower *Int* of the reactivated memory leads to a low Δ*L*, whereas a high *Int* results in an increased degree of Δ*L*. Intriguingly, while the network’s malleability increased with *Int*, we observed a nonlinear response, that is, a distinctive inverted U-shape. This phenomenon was particularly notable when the original network occupied an intermediate modular configuration situated between highly segregated and highly integrated forms, and for intermediate values of *Int*, where Δ*L* showed the greatest sudden increase. Finally, we examined the influence of *θ* on Δ*L* (Figure 3D). From this analysis, it became apparent that *θ* of the network delineated the phase space into two halves, with more stringent *θ* thresholds constraining the spread of activation and consequently diminishing Δ*L*.

In summary, our simulation revealed that after a single reactivation, initial memory stages tend to shift towards a more integrated configuration, and this transition becomes more pronounced with higher reactivation intensities. Conversely, the extent of malleability depends on the interplay between reactivation intensity and the original memory’s degree of integration. Having shown that memory reactivation is effective in inducing structural changes in our memory network model, we next aimed to characterize the evolution of memories over time through repeated reactivations.

##### “Enlightened Stage”: the final state

We investigated how repeated reactivations impacted on the degree of integration (*Z*) in a memory network, simulating 5 reactivations with varied excitability thresholds (*θ*) and intensities (*Int*). Starting with a highly segregated network (*Z*_*o*_ = 0.01) (Figure 4A), diverse speeds of network transformation were observed, with all networks eventually achieving a ceiling integration state (*Z* = 0.76). Notably, the combination of lower *θ* and higher *Int* accelerated integration. Figure 4B and 4C illustrate the dynamics of *Z* for selected *Int* and *θ*, respectively. Higher *Int* led to a faster states of of high integration, reaching it in a single reactivation at the highest *Int*, while increasing *θ* slowed down the transition to high integration states. This emphasizes that *Int* plays a more prevailing role in transforming memory networks from segregated to integrated forms compared to *θ*.

**Figure 4.**
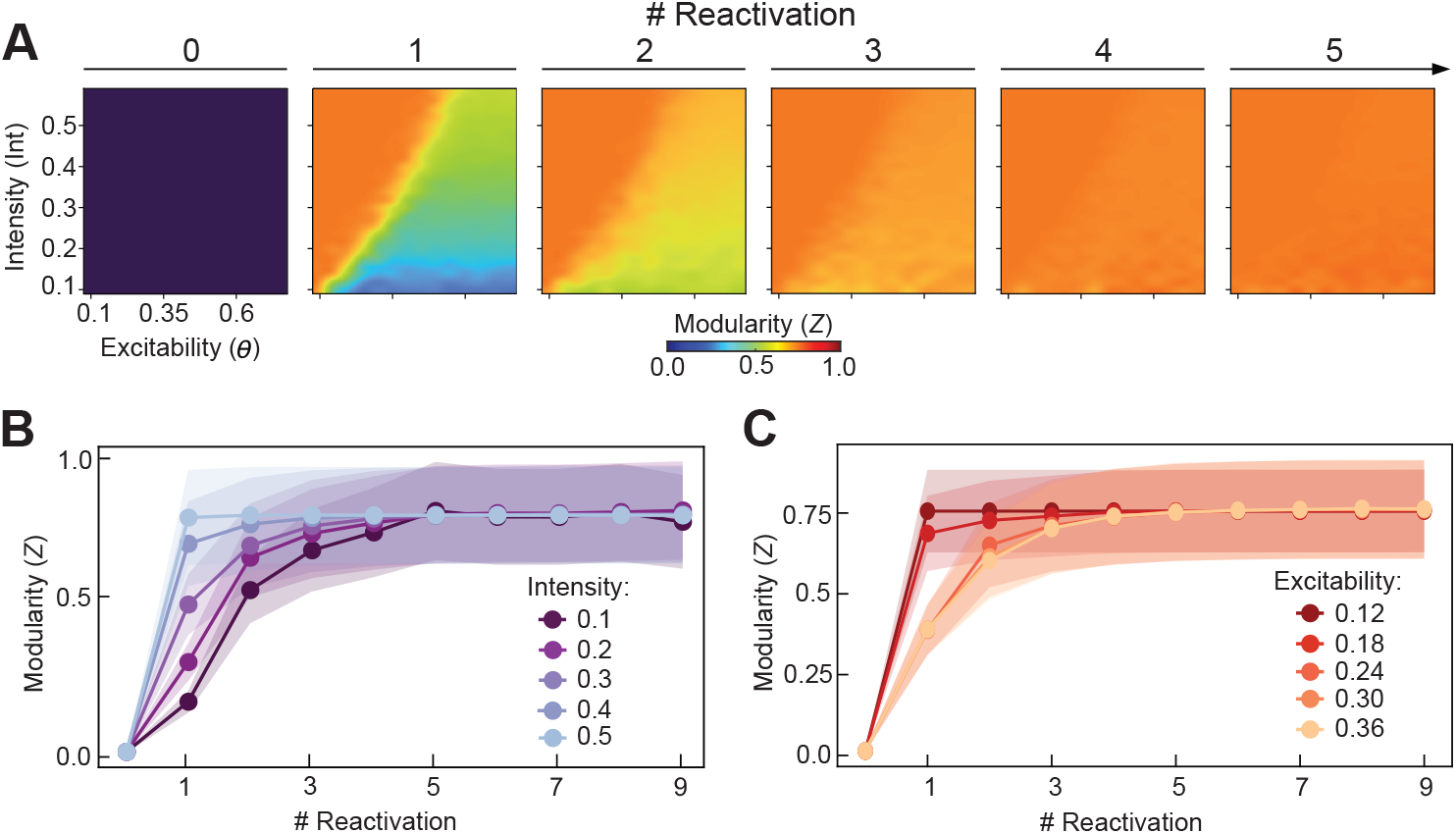
Changes in the degree of modularity of the network for repeated reactivations. **A**. Transformation of *Z* over 5 repeated reactivations. Each panel is a heatmap of *Z* as a function of Intensity and the excitability threshold. After 5 reactivations, the network is completely integrated with *Z*_*o*_ = 0.76. **B**. Curves of *Z* for different intensities and fixed excitability threshold (*θ* = 0.4, mean +- SE). **C**. *Z* transformation for different thresholds (*θ*) and fixed intensity (*Int* = 0.3, mean +- SE). The data reported is the outcome of 25 runs with the same initial state with randomized activated nodes. The initial network has 128 nodes, 4 similar communities of 32 nodes each, and *Z*_*o*_ = 0.01.

Finally, we examined how repeated reactivations impact on the degree of malleability of the network. To address this issue, we quantified the Malleability Index of an initial segregated network form (*Z*_*o*_ = 0.01) as a function of repeated reactivations with varying levels of *Int* and *θ*. These analyses revealed that, for repeated reactivations, the peak of malleability was relocated within the parameter space (Figure 5A). Concretely, under low *θ*, the malleability peak was observed during the initial reactivation, with subsequent reactivations exhibiting minimal changes in their structural configuration. Conversely, more restrictive *θ* led to almost null malleability throughout reactivations, indicating marginal alterations in the number of edges of the network. Interestingly, intermediate levels of *θ* elicited clear malleability peaks observed at the variable reactivation stage, depending on the *Int* of the reactivation. To exemplify the behavior of malleability, we plotted the dynamics of this measure as a function of *Int* for a selected *θ* value (*θ* = 0.4; Figure 5B). These curves displayed revealed important distinctions, particularly in the first reactivation, where the curve corresponding to the highest *Int* exhibited a pronounced maximum malleability. This maximum gradually diminished with increasing *Int*, in line with the observations in Figure 3E. The other curves also indicated peaks of malleability for early reactivations, albeit with lower amplitudes. These findings substantiate previous results when investigating the consequences of a single reactivation, underscoring that memory malleability conforms to an inverted U-shape function, whereby specific windows of opportunity may be optimal for instigating changes in the structure of a previously encoded memory through reactivation.

**Figure 5.**
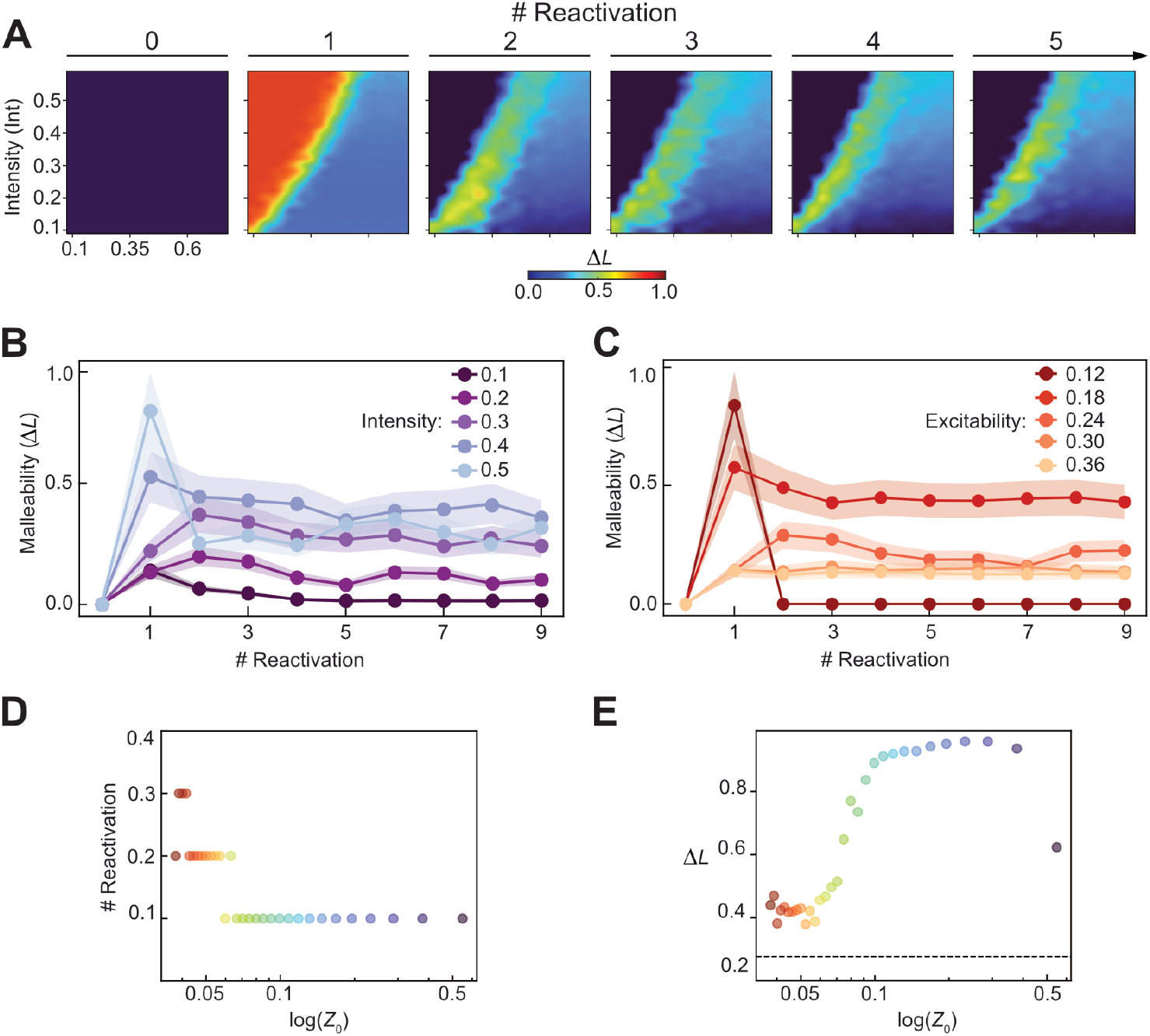
Changes in the Malleability index over repeated reactivations. **A**. Transformation of Δ*L* over 5 repeated reactivations. Each panel is a heatmap of Δ*L* as a function of both intensity and the excitability threshold. The initial state has *Z*_*o*_ = 0.01. **B**. Curves of for different intensities and fixed excitability (*θ* = 0.4, mean +- SE). **C**. Δ*L* dynamics for different excitability thresholds and fixed intensity (*Int* = 0.3, mean +- SE). **D**. Position (in terms of the number of reactivations) of the maximum of Δ*L* as a function of the degree of integration of the initial network configuration (*Int* = 0.3, *θ* = 0.4). **E**. Amplitude of the maximum of Δ*L* as a function of the degree of integration of the initial network (*Int* = 0.3, *θ* = 0.4). The data reported is the outcome of 25 runs that have the same initial configuration with randomized activated nodes. The initial network has 128 nodes, 4 similar communities of 32 nodes each.

Our previous analysis when studying the effects of a single reactivation in the memory network revealed that memory malleability was more sensitive to *θ* than *Int* (e.g., Figure 3B). To further examine this issue in the context of repeated reactivations, we quantified the degree of malleability throughout repeated reactivations with varying degrees of *θ*. The results of these simulations are displayed in Figure 5C. We found that under more permissive *θ* (i.e., *θ <* 0.3), a peak of memory malleability was evident in the initial reactivations, gradually diminishing subsequently. Interestingly, following the attainment of the maximum, malleability persisted, as Δ*L* remained above 0, but at a much lower level, indicating that memories were permeable to structural changes over the entire course of repeated reactivations. In contrast, under more restrictive *θ* conditions (i.e., *θ >* 0.3), malleability remained very low throughout all reactivations, indicating that, under these conditions, memories are minimally susceptible to changing their structure over time.

Together, we found that malleability exhibited a nonlinearity in the context of memory transformation, characterized by an inverted U-shape pattern. This implies that memories within intermediate modular configurations manifest the highest level of malleability. To determine the location and magnitude of the malleability peak, we analyzed the position and amplitude of the nonlinearity for a set of initial networks with varying degrees of integration (*Z*) (with intensity and the excitability threshold fixed, *θ* = 0.4,*Int* = 0.3), as illustrated in Figures 5D and 5E, respectively. Networks with greater segregation peaked later and with a smaller magnitude, while the magnitude of the peak followed a linear increase as the initial network lost its modularity.

## 3 Discussion

The theoretical model presented here, the Segregation-to-Integration Transformation (SIT) model, provides a normative and quantitative framework for assessing the conditions under which memory evolves. The central premise of SIT is that over time, memories shift from highly segregated (i.e., highly modular) to integrated (i.e., less modular) network forms, guided by neural network reactivations, activation spread, and plasticity rules. In addition, our model identifies an optimal window of memory malleability, revealing a nonlinear or inverted U-shaped function for memory evolution. Memories are most malleable at early stages and gradually become less susceptible to changes over time, unifying experimental phenomena and integrating consolidation and reconsolidation accounts. SIT simplifies memory evolution, summarizes it through architectural neural network property rules, and emphasizes the dynamic and optimal malleability of memories throughout the transformation process.

Our model proposes that the transformation of memory relies on modifications in connectivity patterns among nodes within the memory network. This perspective aligns with the idea that learning involves modifications to the wiring diagram of a neural ensemble, where previously unconnected units establish connections and vice versa. However, while alterations in the wiring engram offer a potential substrate for encoding more extensive information in sparse coding models [Knoblauch et al., 2010, Chklovskii et al., 2004], it also underscores that other forms of plasticity based on changes in synaptic weights, the strength of connections between cells, are crucial for comprehending memory evolution [Chklovskii et al., 2004, Bonhoeffer and Yuste, 2002]. Although both mechanisms are likely involved in engram cell formation and function [Poo et al., 2016], investigations of network properties in hippocampal code representation have shown that wiring diagrams effectively encode specific experiences [Ortega-de San Luis et al., 2023, Ryan et al., 2021b]. In addition, shifts in these connectivity patterns provide a more robust explanation for memory transformation than alterations in the individual firing properties of the neurons [Gava et al., 2021]. In subsequent work, the simplicity of the model may enable us to readily incorporate new variables that account for changes in synaptic weights, encompassing both modifications in the wiring diagram and the strength of the connections.

The most notorious finding of the SIT model is that a simple network property, such as modularity of the neural network, leads to a trajectory of memory transformation at the service of generalization [Sun et al., 2023]. In addition, this approach reconciles the theoretical findings in memory consolidation and reconsolidation literature. These models commonly argue that memories are transformed over time; however, while reconsolidation views offer a one-shot direction of the effects, the consolidation view highlights that memories can undergo continuous changes over time [Dudai and Eisenberg, 2004]. The results of our modeling approach offer a reconciling framework by showing that, while memories can evolve perpetually, there may exist an optimal window of malleability during this course. These optimal memory malleability windows appear in the early stages of memory formation, in line with the notion that reconsolidation is a time-dependent phenomenon, as young memories are susceptible to disruption, while older ones are more resistant to change [Milekic and Alberini, 2002, Eisenberg and Dudai, 2004, Suzuki et al., 2004, Frankland and Bontempi, 2005, Alberini, 2011]. Moreover, our findings align better with the idea that, once this malleability opportunity window has passed, memories may still change, albeit at a minimal level [McKenzie and Eichenbaum, 2011, Roüast and Schönauer, 2023].

We conceptualized that memory changes were driven by reactivating the structural properties of a memory network and that the effects of this reactivation would be modulated by the interplay of state-dependent parameters such as the threshold of excitability of the existing network and the intensity of reactivation. The excitability threshold was included in our model to accommodate findings from rodent studies that showed the recruitment of cells to encode memories for a specific event, or engram, hinged on the intrinsic excitability state of the cell, a phenomenon termed memory allocation [Josselyn and Frankland, 2018b]. This line of research provided evidence that fluctuations in neuronal excitability dictate how engrams interact, promoting either memory integration (co-allocation to overlapping engrams) or separation (disallocation to nonoverlapping engrams) [Josselyn and Frankland, 2018b]. On the other hand, the intensity of reactivation was included as a proxy to manipulate the amount of neuron coactivation driven by the overlap between the current input and existing memory [Wammes et al., 2022]. While a pure Hebbian perspective would predict that the greater the degree of coactivation, the larger the degree of integration, recent theoretical accounts, supported by human findings, suggest a more complex relationship between coactivation and representational change than the linear positive relationship predicted by classic Hebbian learning. The nonmonotonic plasticity hypothesis [Detre et al., 2013, Newman and Norman, 2010, Ritvo et al., 2019] proposed a ‘U-shaped’ pattern of representational change based on the degree of coactivation between two neural ensembles. Accordingly, low coactivation results in no overlap change, high coactivation strengthens connections and promotes integration, and moderate coactivation leads to differentiation [Hulbert and Norman, 2015, Wimber et al., 2015, Ritvo et al., 2019]. When applying these parameters to our modeling approach, we found that both the degree of intrinsic excitability and intensity were important in determining the rate at which memory would shift from a segregated to an integrated form. More permissive excitability thresholds and higher intensities, increased propagation, allowing more nodes to be “active”, and promoting memory malleability. More stringent excitability thresholds and lower intensities restricted the spreading of activation in the network and decreasing memory malleability. However, intermediate levels of the parameters induced distinct malleability peaks at variable reactivation stages. Notably, the maximum malleability was located in a state in where the memory configuration was not very segregated and was not integrated, in line with the nonmonotonic plasticity hypothesis.

Our memory model aligns with the idea that a modular architecture that allows independent adjustments within memory networks is beneficial for maintaining enriched detailed memory. However, our framework proposes that the breakdown of modularity occurs as memory evolves as a consequence of repeated reactivation, perhaps cued by overlapping novel experiences. We propose that the shift of memories from segregated to integrated forms responds to the adaptation of the memory system to highly dynamic environments, where novel and past experiences may only partially overlap, necessitating a balanced interplay between stored information and accessibility. In this context, highly segregated memories may function sub-optimally owing to their limited accessibility [Hintze and Adami, 2008]. In contrast, highly integrated memory network forms prioritize accessibility over storage quality, offering an adaptive structural configuration in ever-changing contexts. Network science has demonstrated that at the intersection of these two extreme network configurations, there exists an optimal modularity configuration that effectively balances memory storage and information diffusion [Rodriguez et al., 2019, Nematzadeh et al., 2014]. Similarly, we propose that such an optimal modularity structural state of a memory form is ideal for inducing changes because it provides a balanced trade-off between the probability of being reactivated and the diffusion of activation throughout the network.

While the SIT Model provides a low-dimensional framework in which memory is proactively shaped into a generalized form that involves an integrated state, it fails to address important issues. One such open question pertains to the distribution of network configuration and how the network configuration and its transformation are distributed in the brain. Our model posits that these transformations may occur throughout various brain regions; however, existing models suggest that changes in this network may coincide with localized alterations in specific brain regions [Squire, 1992]. However, others have emphasized the distributed nature of memory formation and transformation [Tonegawa et al., 2018]. In addition, the SIT Model does not consider the selection of neurons that participate in memory encoding or memory allocation [Han et al., 2007]. Although our model includes the excitability threshold, which plays a relevant role in engram formation, we only considered the case where all neurons have the same excitability value. Future studies are needed to determine neuronal competition during memory formation. Finally, the SIT Model does not account for the contribution of sleep to memory consolidation. Although sleep is well known to be critical for changes in the connectivity pattern, there is a wide range of theories supported by numerous experimental findings that point to divergent roles of sleep [Rennó-Costa et al., 2019]. On one hand, theories support sleep as responsible for the renormalization of the strength of the synaptic network, which is fundamental for forgetting irrelevant memories [Tononi and Cirelli, 2014]. On the other hand, the repetitive replay of neuronal firing during offline periods enhances the retention of the most valuable information [Brodt et al., 2023]. While we await a better characterization of this issue at the cellular and structural level, our model offers a simple guiding principle based on the topological structure of neural networks by which memories transform over time. We anticipate that the current model will be of interest to psychologists and neurobiologists to find motivation in testing and challenging the SIT model.

## 4 Material and Methods

Simulations were done using homemade Python codes mainly using Networkx (https://networkx.org/, a package for the creation, manipulation, and study of the structure, dynamics, and functions of complex networks).

### 4.1 Building networks

The Segregated-to-Integrated Model studied the evolution of a set of one-layer, unweighted, and undirected networks. We only analyzed the simplest case and analyzed simulations where the initial condition was a network of 128 equal nodes and four equal communities of 32 nodes. We adopted this simplistic approach to thoroughly explore the roles of different parameters. The degree of the initial modularity of these networks was tuned by varying the mean node degree within the community (*k*_*int*_: from 1 to 31) and the number of edges that connected different communities (*L*^*ext*^: from 1 to 128). To quantify the degree of integration, we defined the parameter *Z* (Equation 3). We used the block model approach to build each community [Holland et al., 1983]. We then linked the communities with a number of edges (*L*^*ext*^) that were assigned randomly to pairs of nodes in different communities. This arrangement ensured that every node and community was not isolated and that they played a similar role in the network.

### 4.2 Reactivation

To shift the nodes from inactive to active mode (*turn-on* step), we introduced the reactivation intensity (*Int*) as a parameter ranging between 0 and 1. The number of activated nodes per community was determined by sampling from a normal probability distribution with a mean of *Int* and a standard deviation of 0.05. *Int* ranged from 0.1 and 0.6, with increments of 0.02. To determine which nodes became active, we utilized a uniform random number generator that selected values between 1 and 32.

The primary parameter of propagation is the excitability threshold *θ*. Throughout the simulations, *θ* varied between 0.1 and 0.8, with increments of 0.05. Spreading was modeled using a deterministic linear threshold rule (Equation 1), which implies that once a node becomes active, it will remain forever, and the propagation will continue until the system reaches a steady state or exceeds 50 iterations [Nematzadeh et al., 2014].

Finally, we applied the *plasticity rule* (Equation 2), where edges were rewired. If two nodes achieve an active mode, an edge is created between them. If two nodes remained inactive, the edges between them were removed. Note that the *Plasticity rule* can be expanded to a weighted network by adjusting the edge weights with a fixed positive (negative) parameter between two active (inactive) nodes. This adjustment indicates an increase (decrease) in strength.

### 4.3 Memory evolution

We simulated the evolution of memory networks by applying 10 consecutive reactivations. The intensity (*Int*) remained constant throughout each reactivation. The nodes activated in each iteration were randomly assigned. We repeated the entire network evolution with the same set of parameters 25 times to characterize the main transformation behavior, independently of the random assignment of nodes. To analyze the outcome of only one reactivation (Figure 3), we focused on the first iteration of the sequence of 10 repeated reactivations.

### 4.4 Quantification of memory transformation

We studied changes in the connectivity patterns of the networks printed by reactivation. First, we analyzed the transformations in the modular structure of the network. We inspected the dynamics of *Z* over a set of repeated reactivations. Also, we figured out shifts of *Z*_*o*_ *→ Z*_*f*_ as a consequence of one reactivation (Equation 4):

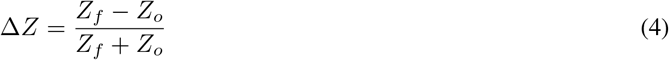

*ΔZ* ranging between -1 and 1; 1 meaning transformation from network to segregated to integrated and -1 the other way around.

Finally, we studied changes in the number of edges. We defined the Malleability Index (Δ*L*) as the number of created (*L*^*c*^)and removed (*L*^*R*^) edges relative to the total number of edges before reactivation (*L*_−1_), Equation 5.

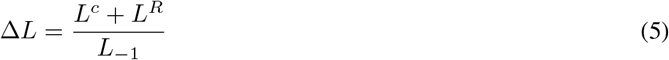

Δ*L* is a positive parameter, 0 value means no change in the number of edges.

## 5 Acknowledgments

This work was supported by a grant from the Agencia Nacional de Promoción Científica y Tecnológica PICT 2020- 00956, to L.B., and by a grant from the Spanish Ministerio de Ciencia, Innovación y Universidades, which is part of Agencia Estatal de Investigación (AEI), through the project PID2022 - 140426NB - I00 (funded by MCIN/ AEI/10.13039/501100011033/ and FEDER a way to make Europe), to L.F. We thank CERCA Programme/Generalitat de Catalunya for institutional support.

